# UNC-45A drives ATP-independent microtubule severing via defect recognition and repair inhibition, contributing to neurite dystrophy

**DOI:** 10.1101/2025.08.26.671351

**Authors:** Asumi Hoshino, Brian Castle, Mihir Shetty, Hannah Khan, Joyce Meints, Olga Pletnikova, Michael K Lee, Juan C. Troncoso, David Odde, Martina Bazzaro

**Author notes:** Corresponding author: Martina Bazzaro Masonic Cancer Center Room 490, 420 Delaware Street S.E Minneapolis, Minnesota 55455.

## Abstract

UNC-45A is the only known ATP-independent microtubule severing protein. Using *in vitro* reconstitution and TIRF microscopy, we show that, unlike canonical severing enzymes such as katanin, spastin, and fidgetin, which hydrolyze ATP to remove tubulin dimers and promote lattice repair, UNC-45A selectively binds to pre-existing microtubule defects and inhibits GTP-tubulin incorporation. This mechanism prevents the formation of stabilized hot spots that typically protect microtubules from disassembly, resulting in persistent lattice damage and net microtubule loss, even in the presence of physiological levels of free GTP-tubulin.

We further demonstrate that UNC-45A localizes near amyloid deposits in both mouse models and human cases of Alzheimer’s disease (AD). In cultured neurons, UNC-45A accumulates in axonal swellings—regions of pronounced microtubule disruption and experimental surrogates for dystrophic neurites in AD—and exacerbates their size and number, particularly under conditions mimicking microtubule damage. Notably, this is the first report of a microtubule severing protein that both localizes near amyloid plaques in tissue and accumulates in neurite swellings in cultured neurons, where it modulates their pathology.

Together, our findings establish the mechanism of ATP-independent, damage-responsive severing pathway that couple defects recognition to repair inhibition, defining a new paradigm in microtubule quality control with broad implications for cytoskeletal integrity and remodeling in health and disease.

## Introduction

Neuronal microtubules are dynamic cytoskeletal polymers that maintain cellular architecture, facilitate intracellular transport, and organize signaling pathways [1, 2]. These microtubules are frequently subjected to structural defects caused by mechanical stress — for example, in curved regions or in regions of their lattice where they contact other microtubules or cellular components — as well as by exposure to damaging agents, including reactive oxygen species [3–10]. Disruption of microtubule integrity is a hallmark of neurodegenerative disorders such as Alzheimer’s disease (AD), where amyloid plaque deposition is associated with local cytoskeletal disorganization, neurite swelling and neuritic dystrophy, and synaptic loss [11–14]. The role of microtubule-stabilizing and protective proteins such as tau and MAP2, in maintaining microtubule stability and protecting neurons from neurodegeneration is well documented [15–19]. Alternatively, while microtubule destabilizing proteins, such as spastin, has been implicated in AD [14], the role of other microtubule destabilizing proteins in neurodegeneration remains untested.

Microtubule severing enzymes cut microtubules along their lattice, causing local changes in the stability and mass of neuronal microtubules. Canonical microtubule severing proteins such as katanin, fidgetin, and spastin are AAA ATPases that use the energy from ATP hydrolysis to remove tubulin subunits from the lattice [20–23]. This creates defects in the microtubule lattice, which can be repaired if free tubulin is available in the system, leading to the formation of regions with a stabilized lattice and a net increase in microtubule mass [24, 25]. By modulating microtubule architecture and interactions with associated proteins such as tau, canonical severing enzymes have been suggested to play a role in neurodegenerative processes [14, 26–28]; however, whether these canonical microtubule severing proteins directly contribute to neurodegeneration remains unclear.

We have recently identified UNC-45A as a novel microtubule-severing protein that, unlike canonical severing enzymes, lacks an ATPase domain yet can directly sever microtubules without ATP hydrolysis [29–34]. UNC-45A is particularly abundant in microtubule-rich neuronal regions of the brain [33] and preferentially binds to curved microtubules exacerbating their bending which results with severing [35]. This unique mechanism led us to hypothesize that UNC-45A might act as a sensor that selectively recognizes and destabilizes structurally compromised microtubules. Here, we show that UNC-45A localizes to amyloid plaques in both AD mouse models and human brain tissue and accumulates at sites of neuritic swelling—regions characterized by disrupted microtubule architecture and are considered an experimental surrogate for the dystrophic neurites seen in AD [36–38]. Significantly, UNC-45A overexpression in neuronal cell models increases the number and size of swellings, indicating that it can exacerbate local microtubule destabilization in vulnerable regions.

To further elucidate its mechanism, we demonstrate using cell-free reconstitution assays that UNC-45A preferentially binds to and severs microtubules containing chemically or mechanically induced lattice defects, in contrast to ATP-dependent severing proteins. Once recruited, UNC-45A remains stably associated with damaged sites, blocking tubulin incorporation and thereby preventing lattice repair. Furthermore, microenvironmental stressors that generate microtubule damage — including oxidative agents and microtubule-targeting drugs — enhance UNC-45A’s recruitment and severing activity. Together, our findings establish UNC-45A as a damage-sensing, ATP-independent microtubule-severing protein that couples lattice defect recognition to persistent destabilization and inhibition of repair. By uncovering this mechanism, we expand the framework for how microtubule networks are remodeled in neurons under stress and provide new insight into the regulation of cytoskeletal integrity in neurodegenerative disease.

## Results

### UNC-45A localizes in the amyloid plaques characteristic of Alzheimer’s disease (AD)

UNC-45A is a microtubule-severing protein recently characterized in our laboratory as the only known ATP-independent microtubule-severing protein [29–34]. Although UNC-45A localizes with microtubules in various cell types and tissues, its expression is particularly elevated in the nervous system, especially in microtubule-rich regions [33]. This spatial pattern suggests a potential role for UNC-45A in regulating neuronal microtubule dynamics. Given the well-documented loss of neuronal microtubule mass and structural degeneration of neurites and synapses in Alzheimer’s disease (AD) [12, 39], we hypothesized that UNC-45A might contribute to microtubule destabilization leading to neuritic dystrophy. To explore this, we analyzed UNC-45A localization relative to amyloid plaques [11] and peri-plaque areas in both the APP/PS1 mouse model of AD [40] and human AD brain tissue. Amyloid plaques and UNC-45A were visualized by immunofluorescence microscopy using antibodies against amyloid-beta [41, 42] and UNC-45A (see **Extended Data Figure 1**). Our results show that UNC-45A is present in regions containing amyloid plaques in both a mouse model of amyloid pathology and in human AD tissue (**Figure 1A–D**). These findings demonstrate for the first time that UNC-45A localizes to amyloid plaques.

**Figure 1.**
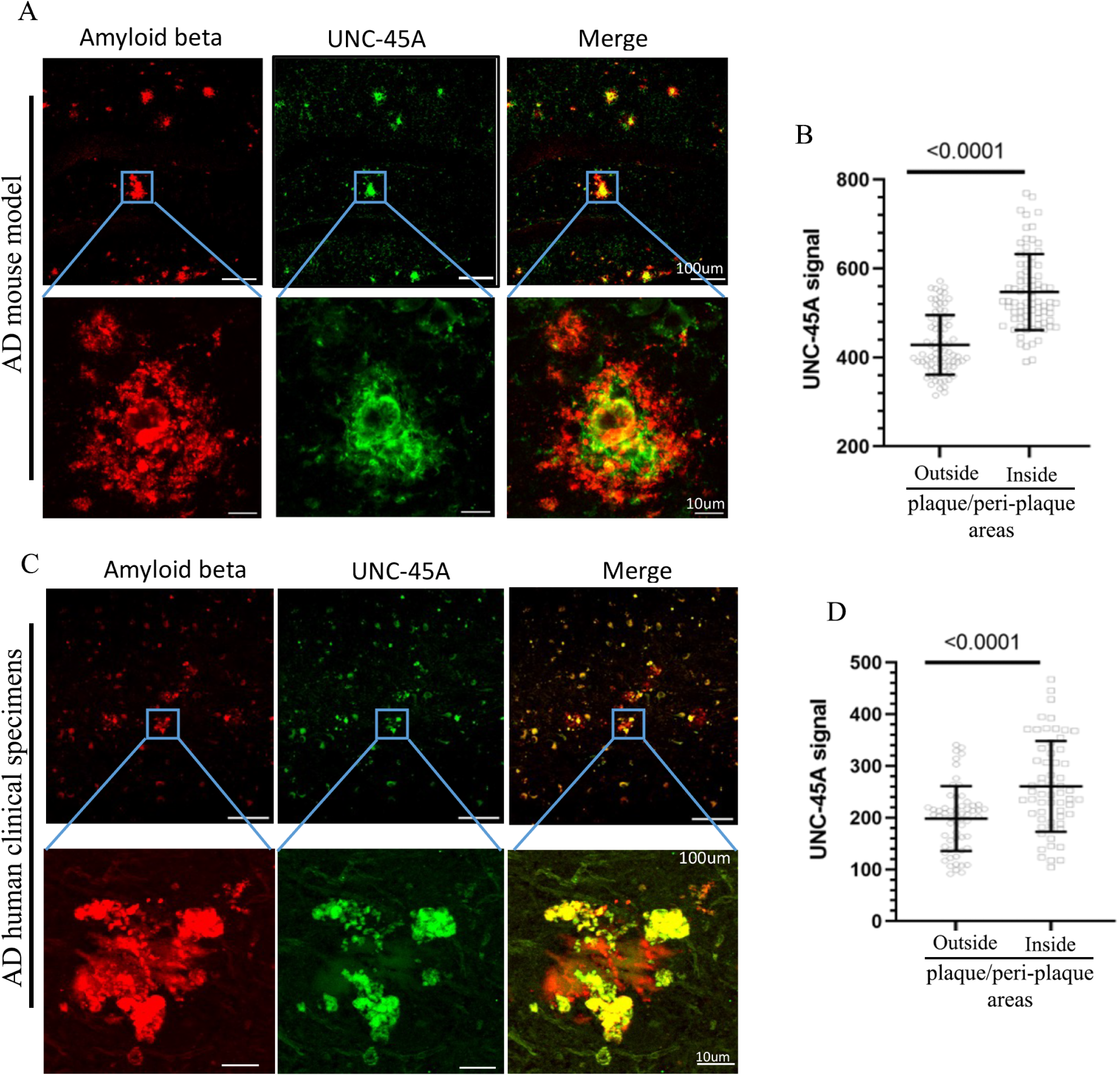
UNC-45A is enriched in the areas containing amyloid plaques in a mouse model of AD and in human specimens of AD. **A**. *Top*, representative images of the dentate gyrus in the hippocampus of APP/PS1 mice co-stained for amyloid beta (red) and UNC-45A (green). The images were captured using a 20x objective. *Bottom*, images show a close-up of the plaque site from the top panels, captured using a 40x objective with maximum intensity projections of Z-stacks. **B**. Quantification of UNC-45A signals outside and inside of plaque and peri-plaque areas in APP/PS1 mouse hippocampi (mean±S.D., n= 80 plaque sites from 4 individual mice per condition). **C**. *Top*, representative images of the dentate gyrus in the hippocampus of human clinical specimens of AD co-stained for amyloid beta (red) and UNC-45A (green). The images were captured using a 20x objective. *Bottom*, images show a close-up of the plaque site from the top panels, captured using a 40x objective with maximum intensity projections of Z-stacks. **D**. Quantification of UNC-45A signals outside and inside of plaque and peri-plaque areas in human clinical specimens of AD (mean±S.D., n= 60 plaque sites from 5 individual clinical specimens).

### UNC-45A localizes at neuronal swelling sites and increases their number and size

Neurite swelling, characterized by disrupted microtubule integrity, are thought to reflect changes associated dystrophic neurites seen in neurodegenerative disorders, including AD [12, 39]. To explore the potential role of UNC-45A in neurite swelling and the microtubule disruption that typically occurs at the site of these swellings, we established primary cultures of hippocampal neurons from newborn mice to examine the subcellular localization of UNC-45A in relation to neuritic swelling. Our findings indicated that UNC-45A is enriched within the swollen regions of growing neurites compared to normal neuritic shaft (**Figure 2A** and **2B**). Analysis of neuronally differentiated mouse neuroblastoma cell line N2A also show enrichment of UNC-45 in the neuritic swelling (**Figure 2C** and **2D**). These findings suggest that UNC-45A may play a role in the formation of these swollen by decreasing microtubule stability in those regions. To determine if UNC-45A expression can regulate neuronal swellings, we overexpressed UNC-45A, via lentiviral transduction, in differentiated N2A cells (**Extended Data Figure 2**) and determined the size and the number of swellings over time. Swelling was defined as a bulge with a diameter greater than twice that of the surrounding neurite [43]. We found that 12 hours post-infection, the number of 2X swelling was similar between the control and UNC-45A overexpression (**Figure 3A**). However, there was a statistically significant difference in the number of swellings when we evaluated both <2X and>2X swelling (**Figure 3B**). After 24 hours post-infection, there was a significant increase in the larger 2X swellings (**Figure 3 C** and **D**) with the noticeable decreased in the smaller swellings. This indicates that the increased expression of UNC-45A increases the number and size of the neuronal swelling in a time-dependent fashion.

**Figure 2.**
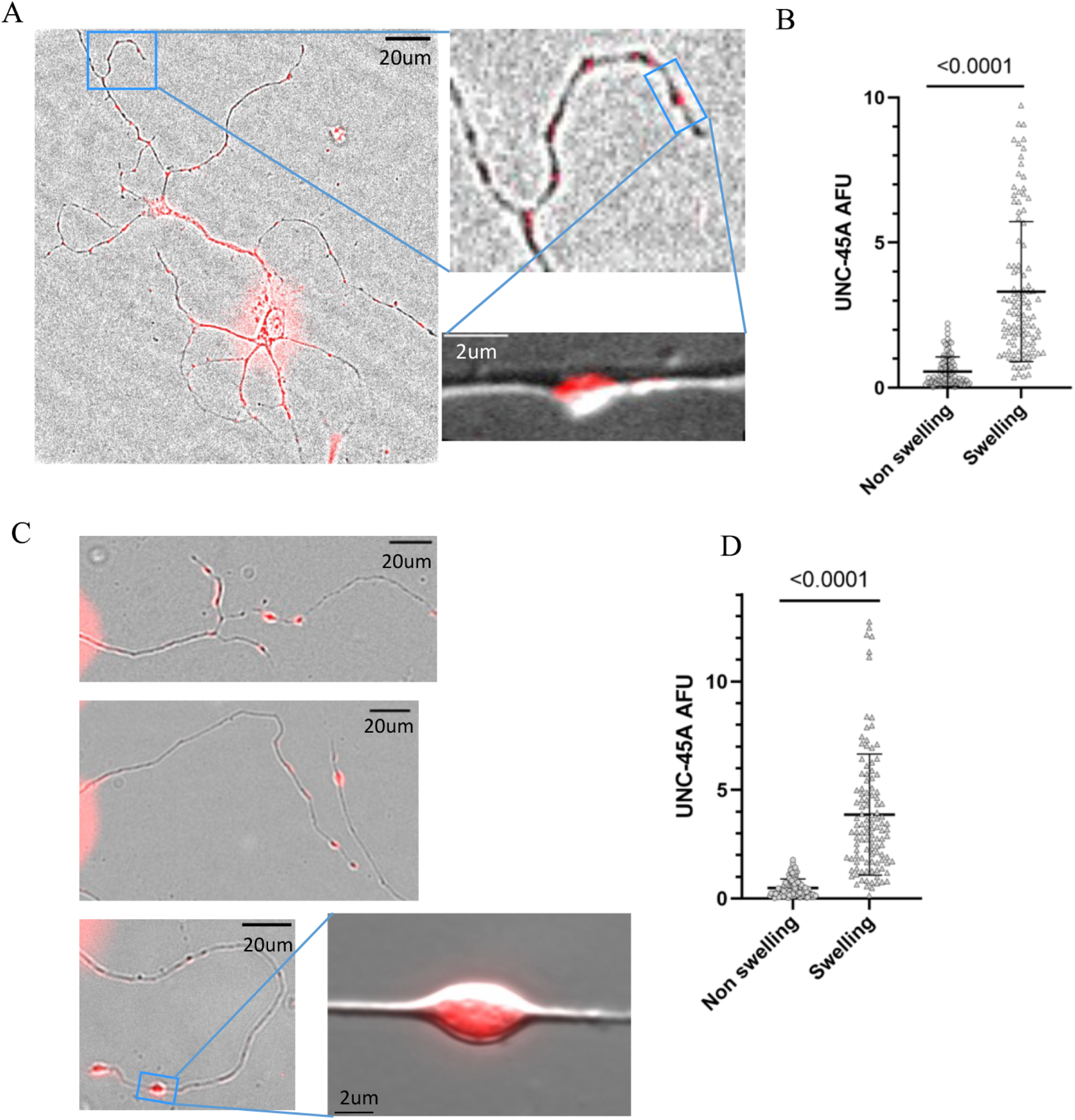
UNC-45A preferentially localizes at the sites of neurite swelling in differentiated neuronal cells. **A.** *Left*, representative image of differentiated mouse hippocampal primary neurons (DIV7) stained for UNC-45A (*red*) and merged with a bright field. The image was acquired with a 40x objective. *Top right*, close-up image of a neurite shown in the *left panel*. *Bottom right,* representative DIC image of neurite swelling shown in the *top right panel*. The image was acquired with a 100x objective. **B**. Quantification of UNC-45A Arbitrary Fluorescent Unit (AFU) at non-swelling and swelling sites shown in A (mean±S.D., n= 102 neurites from 50 individual neurons per each condition). **C**. Representative images (*top, middle and bottom left*) of neurites of differentiated N2A cells (DIV5) stained for UNC-45A (*red*) and merged with a bright field. Images were acquired with a 20x objective. *Bottom right,* representative DIC image of neurite swelling acquired with a 100x objective. **D**. Quantification of UNC-45A AFU at non-swelling and swelling sites shown in C (mean±S.D., n= 104 neurites from 50 individual cells per condition).

**Figure 3.**
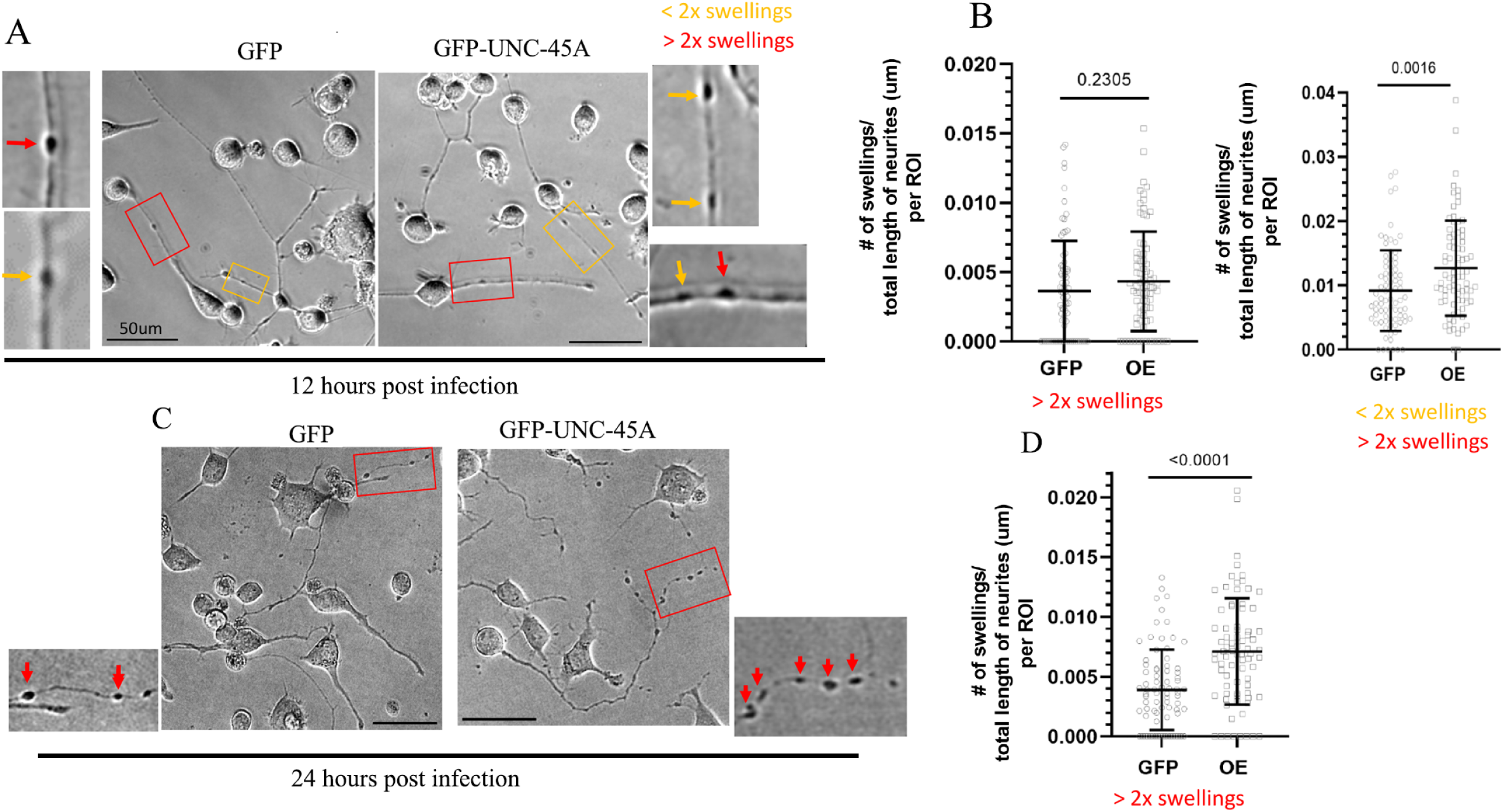
UNC-45A overexpression results in a time-dependent increase in neuronal swelling. **A.** Representative phase contrast images of differentiated N2A (DIV5) expressing GFP or UNC-45A-GFP 12 hours post-infection. *Red boxes* are representative neurite swellings with a diameter greater than twice (>2X) that of the adjacent neurite. *Yellow boxes* represent neurite swellings with a diameter less than twice (<2X) that of the adjacent neurite. Close-ups are also shown for each representative image. **B**. *Left*, Quantification of number of neurite swellings with a diameter greater than twice that of the adjacent neurite (>2X) (mean S.D., n= 80 regions of interest (ROI) per condition, GFP and UNC-45A-GFP overexpressing, OE). *Right,* quantification of neurite swellings including both swellings with a diameter greater and less than twice that of the adjacent neurite (<2X and >2X) (mean S.D., n= 80 ROIs per condition). **C**. Representative bright field images of differentiated N2A (DIV5) expressing GFP or UNC-45A-GFP 24 hours post-infection. *Red boxes* are representative neurite swellings with a diameter greater than twice (>2X) that of the adjacent neurite. Close-ups are also shown for each representative image. Arrows indicate the swelling sites. **D**. Quantification of number of neurite swellings with a diameter greater than twice that of the adjacent neurite (<2X) (mean S.D., n> 80 regions of interest (ROI) per condition, GFP and GFP-UNC-45A overexpressing, OE).

### UNC-45A exacerbates the neuronal swelling caused by microtubule damaging agents

Oxidative stress has been in implicated in the progression of AD [3]. Reactive oxygen species (ROS) have been shown to induce axonal swelling, or beading, in cultured human neurons from forebrain [9]. To establish a system that mimics that, we treated differentiated N2A cells and exposed them to increasing concentrations of H_2_O_2_ and evaluated the effect of this treatment on the neuronal swelling in these cells. Consistent with previous studies done using primary human neurons [9] we found that both H_2_O_2_ exposure increased the number of neuronal swelling in a dose-dependent manner (**Extended Data Figure 3A and B**). We next overexpressed UNC-45A in differentiated N2A cells and tested the effect of its overexpression on neuronal swelling in the presence of different concentrations of H_2_O_2._ For these experiments, we chose to overexpress UNC-45A for 12 hours to reduce the potential toxicity of combining UNC-45A overexpression with H_2_O_2_ treatment. We found that exposure to H_2_O_2_ led to a significant increase in neuronal swelling in cells overexpressing UNC-45A compared to the control (**Figure 4A** and **B**). In cultured mouse neurons, the microtubule-stabilizing agent taxol has been shown to induce reactive oxygen species (ROS) by increasing NADPH oxidase activity [44]. Therefore, we investigated whether exposure to taxol would influence neuronal swelling in our system. We found that taxol exposure resulted in a significant increase in neuronal swelling in a dose-dependent manner (see **Figures 4C** and **4D**). Additionally, we overexpressed UNC-45A in differentiated N2A cells to examine the effect of its overexpression on neuronal swelling in the presence of varying concentrations of taxol. The results showed that taxol exposure significantly increased neuronal swelling in cells overexpressing UNC-45A compared to the control group (see **Figures 4E** and **4F**). Together, this suggests that UNC-45A plays a role in the development and size of neuronal swelling, and that the presence of microenvironmental stimuli, including H_2_O_2_ or taxol, in the system.

**Figure 4.**
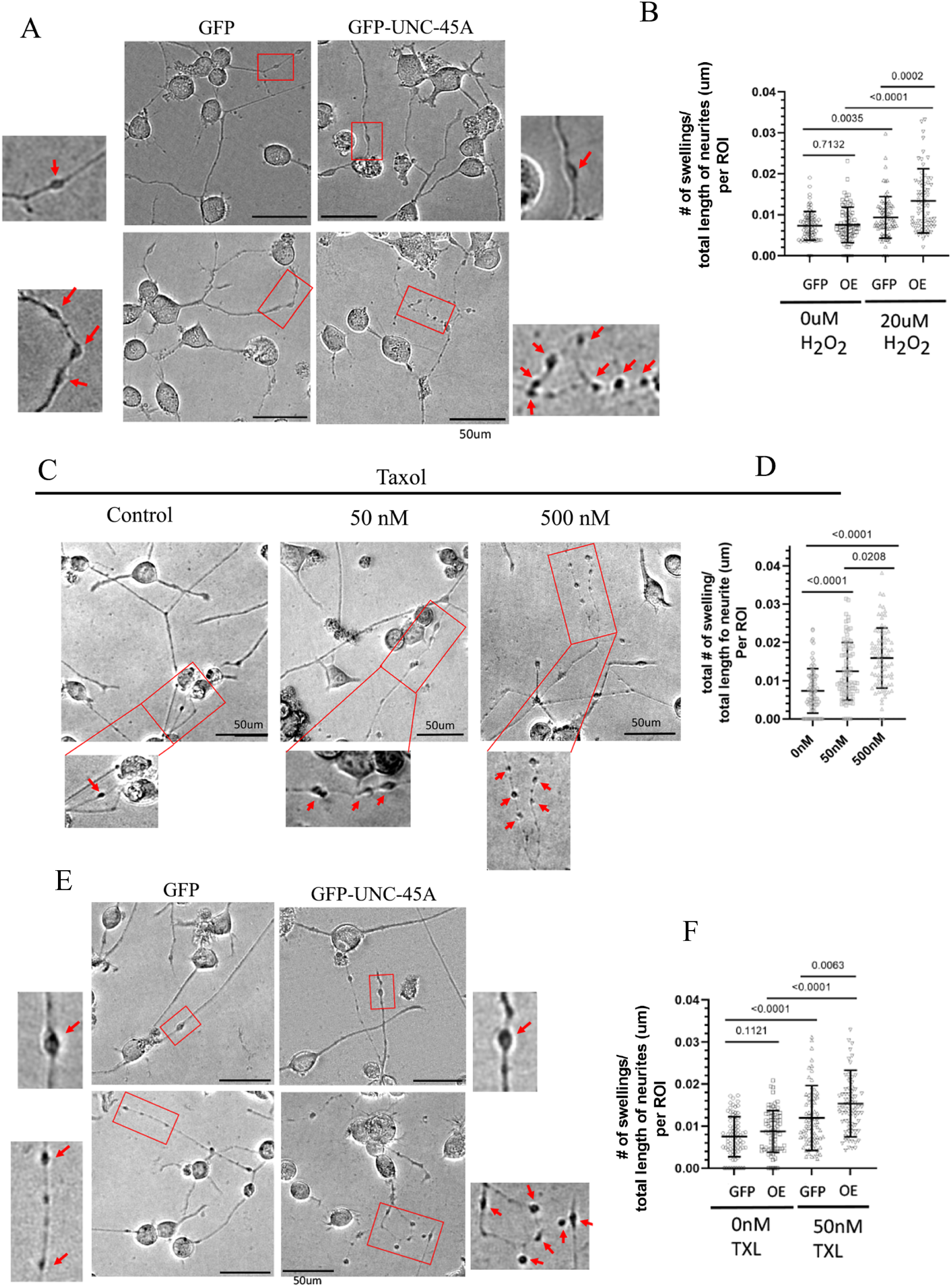
Exposure to H_2_0_2_ and taxol worsens UNC-45A’s impact on neuronal swelling. **A.** Representative bright field images of differentiated N2A (DIV5) expressing GFP or UNC-45A-GFP (12 hours of post-infection) treated with vehicle (*top panels*) or 20𝜇M H_2_O_2_ (*bottom panels*). *Red boxes,* representative neurite swellings with a diameter greater than twice that of the adjacent neurite (>2X). Close-ups are also shown for each representative image. Arrows indicate the swelling sites. **B**. Quantification of number of neurite swellings (mean±S.D., n= 80 ROIs per condition, GFP and UNC-45A-GFP overexpressing, OE). **C**. Representative bright field images of differentiated N2A (DIV5) treated or not with taxol at the indicated concentrations. *Red boxes,* representative neurite swellings with a diameter greater than twice that of the adjacent neurite (>2X). Close-ups are also shown for each representative image. Arrows indicate the swelling sites. **D**. Quantification of number of neurite swellings (mean±S.D., n= 80 ROIs per condition). **E**. Representative bright field images of differentiated N2A (DIV5) expressing GFP (GFP) or UNC-45A-GFP (OE) 12 hours of post-infection treated with vehicle (*top panels*) or 50 nM of taxol. **F**. Quantification of number of neurite swellings. (mean±S.D., n= 80 ROIs per condition, GFP and GFP-UNC-45A overexpressing, OE).

### UNC-45A preferentially binds to and act on damaged microtubules

Exposure of microtubules to H_2_O_2_ or taxol causes chemically induced defects, or nanodamages, to their lattice [45, 46]. Thus, we hypothesized that UNC-45A preferential binds to and acts on damaged microtubules which could explain the increase in neuronal swelling in UNC-45A overexpressing cells when exposed to H_2_O_2_-or taxol. To test our hypothesis, we switched to cell-free reconstitution experiments which enable us to directly assess the interaction and impact of a MAP at the individual microtubule level in a controlled system [30, 46, 47]. We pretreated rhodamine-labeled microtubules with either 200 uM of H_2_O_2_ overnight or 10 or 100 uM of taxol for 1 hour, or their respective vehicle (H_2_O and DMSO respectively). (**Figure 5A**). This concentration and duration of exposure have been previously shown to be sufficient to cause nanodefects in the microtubule lattice and to mimic the intracellular levels these agents would reach in an oxidative environment or following taxol treatment [29, 31, 46]. After agents’ wash out, recombinant GFP-UNC-45A was introduced in the system and its binding to microtubules was evaluated as we previously described [29, 30, 47]. We found that UNC-45A preferentially binds to H_2_O_2_-and taxol-treated microtubules (**Figure 5B**) as indicated by the increase in the UNC-45A signal along treated versus untreated microtubules (**Figure 5C**). In addition to chemically induced nanodefects, microtubule also have mechanically induced defects. These defects can occur when microtubules come into contact with each other in overcrowded areas of the cells, such as during neuronal swelling [48, 49]. Thus, we tested whether UNC-45A also preferentially binds to microtubule in their crossing or bundling areas and found that UNC-45A’ signal increases in crossing and bundling areas of microtubules (**Extended Data Figure 4A and B**). Consistent with our previous findings, the engagement of UNC-54A with microtubule results in their severing in a time and concentration-dependent manner (Figure **5** **D** and **E**).

**Figure 5.**
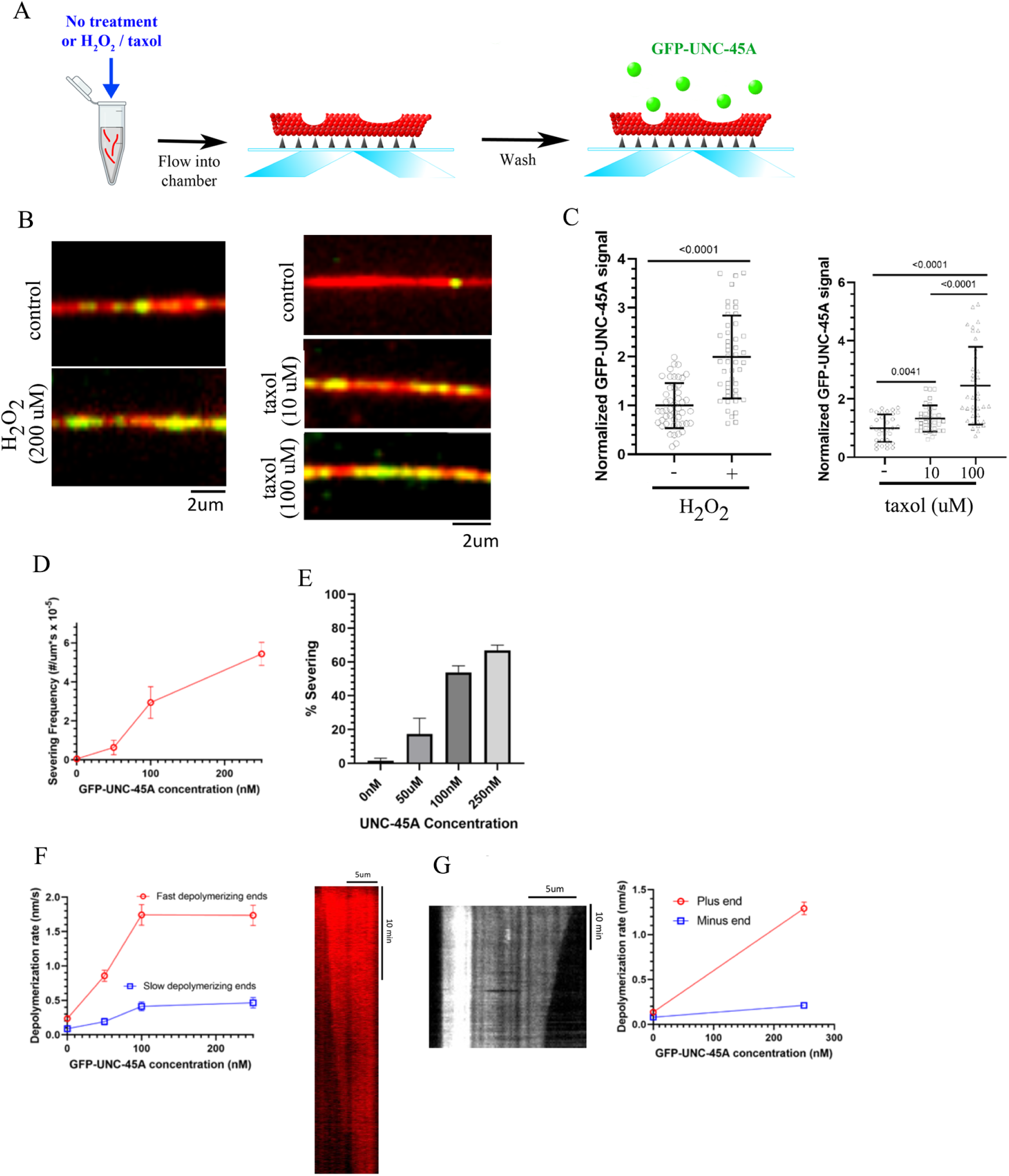
UNC-45A preferentially binds to and acts on defective microtubules. **A.** Experimental set-up for microtubule-defect experiments. **B.** *Left*, representative images of GFP-UNC-45A (green) binding to rhodamine-labeled GMPCPP microtubules (red) treated or not (vehicle, water) with H_2_O_2_ (200µM overnight). *Right*, representative images of GFP-UNC-45A (green) binding to GMPCPP microtubules (red) treated or not (vehicle, DMSO) with taxol (0, 10, 100 µM for 1hr). **C**. *Left*, quantification of GFP-UNC-45A binding to microtubules treated ± H_2_O_2_. The fluorescent intensity of GFP-UNC-45A on single microtubules in each condition was quantified (mean±S.D., n= 48 microtubules per condition). *Right*, quantification of GFP-UNC-45A binding to microtubules treated with taxol. The fluorescent intensity of GFP-UNC-45A on single microtubules in each condition was quantified (mean±S.D., n= 32 microtubules for 0μM, 42 microtubules for 10μM, and 38 microtubules for 100μM taxol). **D**. Severing frequency in the presence of 0 nM (n=130); 50 nM (n=55); 100 nM (n=68); 250 nM (n=189) GFP-UNC-45A. n-value represents the number of microtubules analyzed, the points on the plot represent the mean value, and the error bars represent the mean ± SE. **E**. Percentages of microtubules that displayed at least one resolvable severing event during the microtubule-severing assays. For all data displayed, the number of microtubules analyzed is: 0 nM (n=130); 50 nM (n=55); 100 nM (n=68); 250 nM (n=189) GFP-UNC-45A. Error bars represent mean ± SE. **F**. *Left*, quantitative measurement of microtubule depolymerization rate plotted over various concentrations of GFP-UNC-45A (0nM-250nM). The number of microtubule ends analyzed is; 0 nM (n=60); 50 nM (n=67); 100 nM (n=73); 250 nM UNC-45A-GFP (n=75). Error bars represent mean ± SE. *Right*, the representative kymograph showing depolymerization of rhodamine-labeled GMPCPP microtubule treated 10µM taxol in the presence of 250nM GFP-UNC-45A. The kymograph was used to measure depolymerization rates by drawing a line at both ends of the microtubule and measuring the change in distance over the change in time. **G**. *Left*, the representative kymograph showing depolymerization of polarity-marked GMPCPP microtubule treated 10µM taxol in the presence of 250nM GFP-UNC-45A. The bright segment is a GMPCPP seed with a minus end. The dim segment is an elongated microtubule from the seed with a plus end. *Right*, quantitative measurement of microtubule depolymerization rates at minus/plus ends in the absence of GFP-UNC-45A or the presence of 250nM GFP-UNC-45A. The number of microtubule ends analyzed is 0 nM (n=22 for both minus/plus ends); 250 nM UNC-45A-GFP (n=49 for both minus/plus ends). Error bars represent mean ± SE.

To further support the evidence that UNC-45A preferentially acts on defective regions of microtubules, we first investigated whether UNC-45A affects the depolymerization rate differently at the two microtubule ends, and then determined which end — plus or minus — is more affected, given that the plus end is more open and thus more ‘damaged’ than the minus end [50, 51]. For this, we exposed polarity marked microtubules to increasing concentrations of UNC-45A and found that, in the presence of UNC-45A the microtubule plus is 1.29±0.0701 nm/sec *vs*. 0.137±0.0185 in control, while the one of the minus end is 0.151±0.0263 nm/sec *vs*. 0.078±0.0119 in control (**Figure 5F and G**). Together this indicates that UNC-45A is a potent microtubule severing protein that preferentially acts on damaged areas of the microtubules.

### UNC-45A dwells on defective microtubules and prevents their repair

Sites of damage in the microtubule lattice can be repaired by the incorporation of free tubulin, forming so-called ‘hot spots’ [24, 25]. Our data show that exposure to UNC-45A leads to rapid severing of defective microtubules. Here, we wanted to test whether this effect is due to UNC-45A preventing microtubule repair. We first compared how long UNC-45A remains bound to defective microtubules with the behavior of the classical ATP-dependent microtubule-severing protein katanin, which binds and detaches quickly from the microtubule lattice. This transient binding by katanin creates defects but also allows repair to occur. We found that the duration of association for UNC-45A was 305.6 seconds, compared to 26.5 seconds for katanin, as previously reported [52] and experimentally confirmed here. Additionally, the binding frequency of UNC-45A was 1,260 µm⁻¹·s⁻¹, whereas katanin exhibited a higher binding frequency of 4,314 µm⁻¹·s⁻¹ (**Figure 6A** and **B**). This suggests that when UNC-45A is bound to defective microtubules, the addition of free GTP does not lead to its incorporation into the microtubule lattice. To directly test this, we employed an *in vitro* dynamic system in which lattice defects can be repaired in the presence of free GTP-tubulin. Using this system, we first confirmed that microtubules damaged by taxol or H₂O₂ treatment undergo repair when free tubulin is available, forming GTP-tubulin hotspots in an agent-and dose-dependent manner (**Extended Data Figure 5A, B**). Having validated the system, we next exposed taxol-treated microtubules to UNC-45A (or vehicle control), washed out unbound protein, and then introduced free GTP-tubulin. We quantified its incorporation into the microtubule lattice. As shown in **Figure 6C**, in the absence of UNC-45A, free tubulin was readily incorporated. However, when UNC-45A was present, GTP-tubulin incorporation was significantly reduced, reaching levels similar to those of microtubules that were not treated with taxol and therefore contain fewer lattice defects (**Extended Data Figure 6C**).

**Figure 6.**
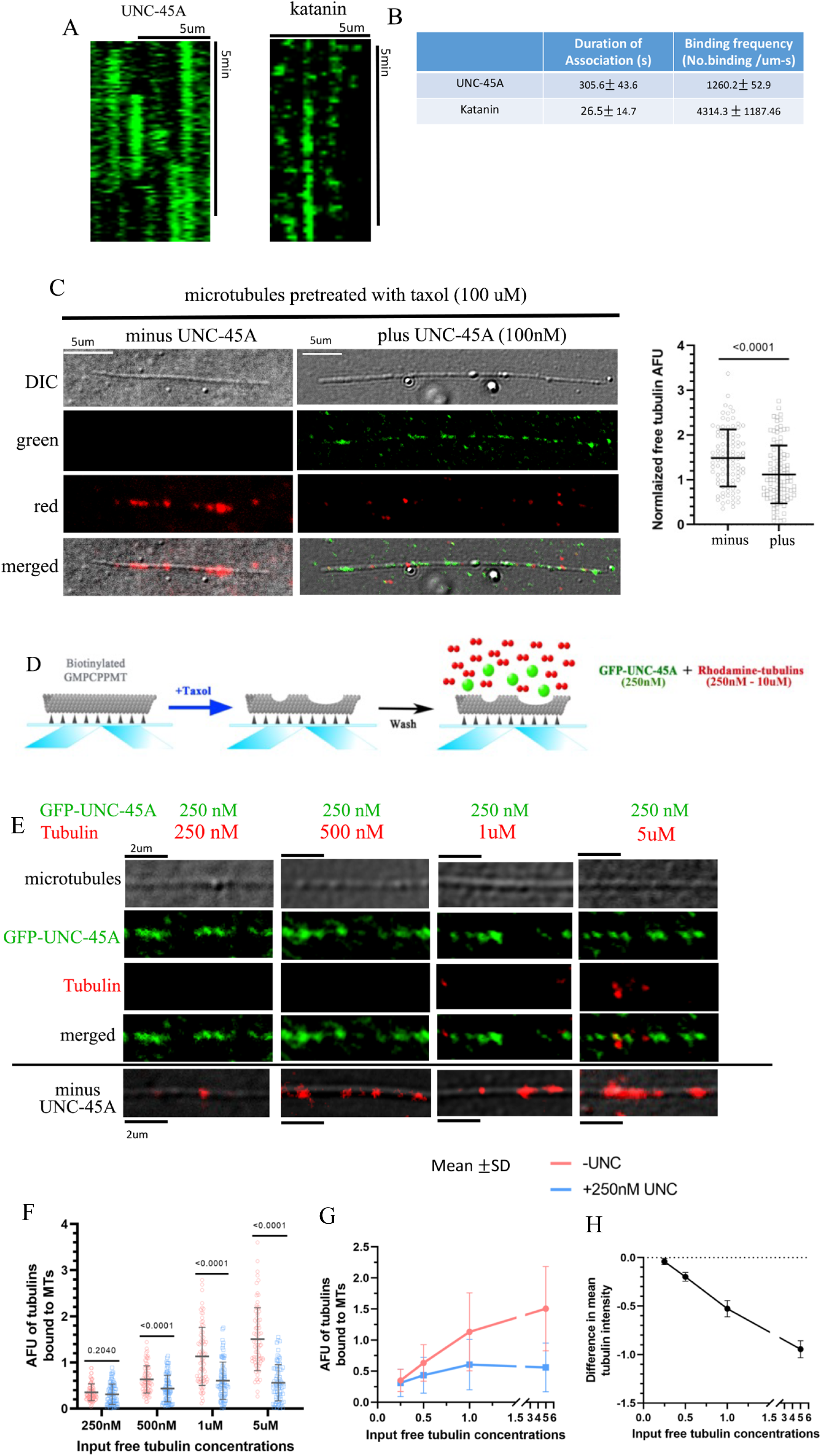
UNC-45A dwells on defective microtubules and prevents their repair. **A.** Representative kymographs showing GFP-UNC-45A and GFP-katanin binding to microtubules. 100nM GFP-UNC-45A or 25nM GFP-katanin were used for the binding assay. These kymographs were used to calculate the duration of association and binding frequency. **B.** Quantification of binding kinetics of GFP-UNC-45A and GFP-katanin to microtubules. N= 527 and 542 binding events were quantified for GFP-UNC-45A and GFP-katanin, respectively. **C.** *Left*, representative images of biotinylated MT (visualized with DIC) treated with 100µM taxol for 1 hr and repaired with 500nM rhodamine-tubulins (red) minus/plus 100nM GFP-UNC-45A (green). *Right*, quantification of fluorescent intensity of rhodamine-tubulins binding to MTs minus/plus UNC-45A and expressed as normalized free tubulin AFU. n= 100 MTs per condition were quantified. Error bars represent mean ± SD. **D.** Experimental set-up for microtubule-repair experiments in the presence of free tubulin and UNC-45A. **E**. *Top four panels*, representative images of biotinylated MT treated with 100µM taxol for 1 hr and exposed (or not, last four panels from the bottom-minus UNC-45A) to 250nM GFP-UNC-45A and increasing concentrations (250nM-5uM) of rhodamine-tubulin. *Second four panels from top*, green channel (UNC-45A). *Third four panels from top,* red channel (rhodamine tubulin). *Fourth four panels from top,* merged image of green and red channels. **F.** Quantification of fluorescent intensity of rhodamine-tubulins binding to microtubules expressed as AFU of tubulins bound to microtubules for different input of rhodamine tubulin. n= 80 MTs per condition from two independent experiments were quantified. Error bars represent mean ± SD. **G**. Mean ± SD of AFU of tubulins bound to microtubules is plotted over the input free tubulin. Error bars represent mean ± SD. **H**. Difference in mean rhodamine-tubulin AFU between minus/plus 250nM GFP-UNC-45A is plotted over the input free tubulin concentrations. n= 80 microtubules per condition were quantified. Error bars represent mean ± SE.

**Figure 7.**
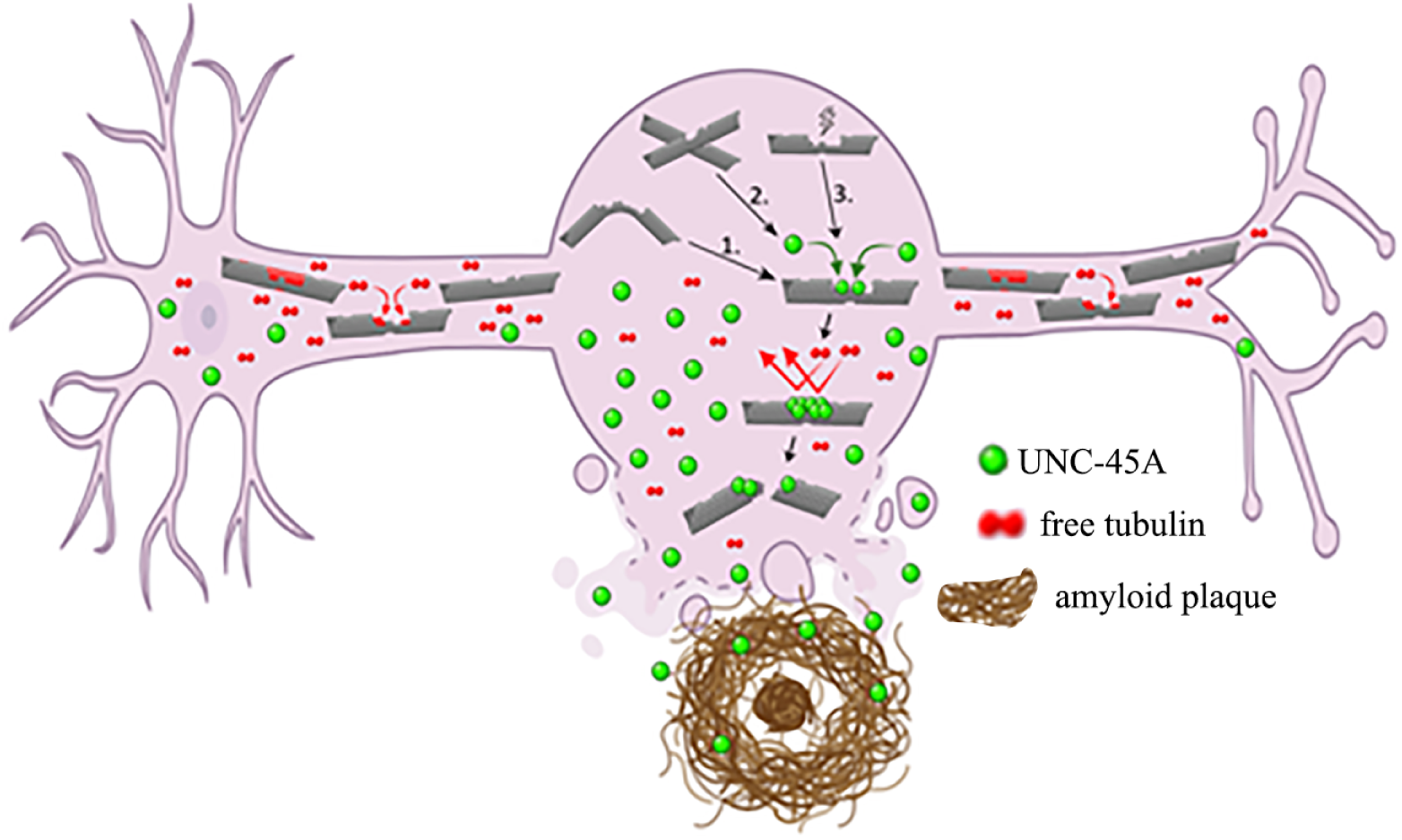
Summary model. Within non-swelling neurites, microtubules undergo occasional damage that is repaired by the incorporation of free tubulin (red), resulting in the formation of GTP-tubulin hotspots. In contrast, at swelling sites (bulges along neurites), microtubules experience severe damage caused by mechanical factors (1. microtubule bending/curving, 2. microtubule crossing) and 3. chemical stress (e.g., H₂O₂). At these sites, the high local abundance of UNC-45A (green) allows it to preferentially bind to damaged microtubule regions, blocking access for free tubulin and preventing repair. This persistent inhibition of repair promotes microtubule breakage and loss, which is typical of amyloid plaques.

Next this dynamic system was used to evaluate the formation of GTP-tubulin hot spots in the presence of both UNC-45A and free tubulin (**Figure 6D**). As shown in **Figure 6E**, when UNC-45A (green) and free tubulin (red)—at concentrations up to 20-fold higher—are simultaneously present near damaged microtubules (visible by phase contrast), UNC-45A binds to microtubules. It strongly reduces the formation of hot spots (**Figure 6E**), in a manner that is dependent on the free tubulin concentration (**Figure 6F** and **G**). We confirmed that the reduction in the free tubulin incorporation in the lattice is not due to free tubulin sequestration or self-assembly, as shown by pellet assays (**Extended Data Figure 5D**). These findings support the conclusion that UNC-45A interferes with the repair of microtubule lattice defects by binding to damaged microtubules and blocking the incorporation of free tubulin at repair sites.

## Discussion

In this study, we show that UNC-45A accumulates in amyloid plaque–rich regions of Alzheimer’s disease brain tissue, which are characterized by local microtubule loss and neuritic dystrophy [11–14]. Using primary hippocampal neurons and differentiated N2A cells, we further demonstrate that UNC-45A localizes to neuritic swellings and that its overexpression increases swelling number and size, particularly under microtubule-damaging conditions. These observations indicate that UNC-45A is not only recruited to sites of damage but also actively contributes to microtubule destabilization in vulnerable neuronal regions.

Through cell-free reconstitution, we reveal that UNC-45A preferentially binds to microtubules with lattice defects, blocks tubulin incorporation, and prevents repair independently of ATP hydrolysis. UNC-45A is the only known non-canonical microtubule-severing protein that does not require ATPase activity [29–34], and to our knowledge, this is the first report of any microtubule-severing enzyme being present at amyloid plaques and enriched in neuronal swellings. Given that ATP depletion is a common feature of neurodegenerative diseases including AD [53] the ATP-independent activity of UNC-45A may confer a functional advantage in sustaining local microtubule destabilization under energy-limiting conditions. Together, these findings identify UNC-45A as a damage-responsive, ATP-independent severing factor that couples defect recognition to persistent destabilization, providing new insight into the regulation of cytoskeletal integrity in neurodegenerative disease.

Comparison with canonical severing enzymes further highlights the novelty of UNC-45A’s mechanism. ATPase-driven severing enzymes not only creates new microtubule ends via the severing process but also defects in the microtubule lattice which, when free tubulin is available, can be repaired, forming stable GTP-rich patches within the lattice that can increase overall microtubule mass and resistance to disassembly [24, 25]. Thus, canonical severing can paradoxically promote increase of microtubule mass, depending on the cellular context. Canonical microtubule-severing proteins are proposed to play pivotal roles in controlling microtubule dynamics during neurodegeneration. In hereditary spastic paraplegia, loss of spastin function results in insufficient severing, which disrupts microtubule dynamics and leads to axonal degeneration [26]. Figetin has been proposed to promote Wallerian-like degeneration specifically in dendrites by actively disassembling microtubules after injury and its knockdown has been shown to promote regeneration [54, 55]. In Alzheimer’s disease, severing proteins have been implicated through their ability to modulate tau and to be regulated by it. This includes spastin’s role in severing microtubules in dendrites of primary neurons exposed to Aβ oligomers, where tau becomes mislocalized [14], and the role of tau in protecting microtubules from severing by katanin by condensing on microtubules to form a barrier that prevents katanin from acting [27, 28].

In contrast, UNC-45A’s ATP-independent activity does not generate repairable defects but instead locks damaged regions into a persistently destabilized state by blocking tubulin incorporation. This mechanism implies that once UNC-45A is recruited to a defective site, that region becomes resistant to normal lattice repair, potentially driving irreversible local breakdown of microtubule architecture. Thus, although in principle UNC-45A–mediated removal of defective lattice segments might help maintain microtubule network fidelity, our data suggest that under chronic stress conditions UNC-45A activity becomes maladaptive, driving persistent destabilization and contributing to net microtubule loss. Furthermore, given the high metabolic demands of neurons and their reliance on stable microtubule tracks for cargo transport and synaptic maintenance, this persistent severing mechanism may represent a double-edged sword for neuronal homeostasis.

An important question to explore is the precise structural basis by which UNC-45A recognizes lattice defects and the characteristics of these defects. It is reasonable to hypothesize that UNC-45A may form oligomers that bind to the defective regions, thereby both inhibiting repair processes and exerting mechanical forces that contribute to lattice destabilization. This potential for oligomerization aligns with findings that the worm isoform of UNC-45A forms oligomeric species [56]. Future high-resolution structural studies will be critical to elucidate these mechanisms. Such insights could uncover novel strategies for pharmacological modulation of UNC-45A activity. Another key question is whether UNC-45A’s severing function could be therapeutically targeted to restore microtubule integrity in AD and related disorders. If UNC-45A-driven destabilization proves to be maladaptive under chronic stress, pharmacological inhibition of its recruitment, oligomerization and severing activity might help preserve microtubule mass, maintain axonal and dendritic architecture, and protect synaptic connectivity.

In summary, our study uncovers a unique, damage-sensing mechanism of microtubule destabilization driven by UNC-45A that operates independently of ATP hydrolysis and preferentially targets lattice defects. By linking defect recognition to persistent local destabilization, UNC-45A expands the paradigm of microtubule quality control and highlights an underexplored layer of cytoskeletal regulation in neurodegenerative disease. Our findings reveal an ATP-independent microtubule damage amplifier, challenging the paradigm that severing must always couple with repair to maintain network integrity.

## Materials and methods

### Antibody and chemicals

Mouse monoclonal anti-UNC-45A (Enzo, ADI-SRA-1800-D) was used for immunofluorescence staining and western blot to detect UNC-45A. Rabbit monoclonal anti-amyloid β (Invitrogen, 700254) was used for immunofluorescence staining. This antibody is well-characterized and specifically recognizes amyloid β (1-42) [57, 58]. Secondary antibodies used were peroxidase-linked anti-mouse IgG (Cell signaling, 7076S), Alexa Fluor 488-conjugated donkey anti-mouse IgG, Alexa Fluor 594-conjugated donkey anti-rabbit IgG, and Alexa Fluor 594 donkey anti-mouse (all Jackson ImmunoResearch Laboratories, 715-545-150, 711-585-152, and 715-585-151, respectively). Taxol was purchased from Cytoskeleton. H_2_O_2_ was purchased from Sigma.

### Mice

APP/PS1 transgenic mice [B6.Cg-Tg(APPswe,PSEN1dE9)85Dbo/Mmjax; 034829-JAX] and age matched controls were used in this study. These mice show progressive amyloid pathology starting at ∼6 mos of age [59]. The mice at 15 months of age were sacrificed by intracardial perfusion of 4% paraformaldehyde in PBS and brain was collected. The brains were processed for immunohistochemistry as previously described [60, 61]. Briefly, the brains were fixed in 4% paraformaldehyde and cryoprotected in 30% Sucrose/PBS solution and coronally sectioned (40 μm) on a freezing sliding microtome (Leica). All experimental protocols involving mice were in strict adherence to the NIH Animal Care and Guidelines and were approved by the Institutional Animal Care and Use Committee (IACUC) at the University of Minnesota. All applicable ethical standards required by University of Minnesota IACUC was followed.

### Human clinical specimens of AD

Human postmortem formalin-fixed paraffin-embedded (FFPE) tissue sections were dissected from the postcentral gyrus of Alzheimer’s disease cases with high level (n=12) and intermediate level (n=4) neuropathologic change [62], and age-matched controls (n=14). These tissues were provided by the Johns Hopkins Brain Resource Center from autopsies of participants in the JHU Alzheimer’s Disease Research Center under protocol approved by the institutional IRB.

### Tissue immunofluorescence Mouse tissues

Immunofluorescence staining of 40-μm coronal floating brain sections was performed as previously described [15, 63] with modifications to improve signal-to-noise ratio. Briefly, sections were rinsed with TBST and subjected to antigen retrieval for 30 min at 75℃ in 1X Reveal Decloaker (Biocare Medical) and 3% hydrogen peroxide for 10 min to reduce autofluorescence. Subsequently, the sections were blocked with Background Sniper (Biocare Medical) for 13 min, and incubated with anti-UNC-45A (1:50 dilution) and anti-amyloid β (1:1000 dilution) in 5% Background Sniper overnight at 4℃. On the following day, the sections were incubated with anti-mouse Alexa 488 and anti-rabbit Alexa 594 secondary antibodies (1:250 and 1:1000 dilutions, respectively) in 5% Background Sniper for 2 hrs at room temperature. The sections were stained with 4′,6-diamidino-2-phenylindole (DAPI) for 15 min and mounted with VectaShield Vibrance antifade mounting medium (Vector Laboratory). Images were acquired with an Olympus Fluoview FV1000 BX2 upright confocal microscope with UPlan S Apo 20x/0.85 NA objective, UPlan FL N 40x/1.30 NA objective, and FluoView software (Olympus) and ProgRes C3 CCD digital camera (Jenoptik). 40x Z-stack images were acquired with a step size of 0.48um and the zooming factor of 4.9 resulting in a pixel size of 0.127μm. Alexa 488 and 594 were excited with 488nm and 543nm lasers, respectively. Images were taken with sequential excitation to avoid bleed-through. Four mice brains were examined for UNC-45A accumulation in amyloid plaques.

### Human Tissues

Immunofluorescence staining of 5-μm paraffin embedded fixed brain slides was performed similarly to mouse tissue immunofluorescence except for deparaffinization and antigen retrieval. Tissue slides were deparaffinized with 100% xylene and rehydrated with gradient ethanol (100%, 95%, and 80%). Antigen retrieval was then carried out with 1X Diva Decloaker (Biocare Medical) in a vegetable steamer for 30 min at 100℃. Images were acquired with the same microscope setting as mouse tissue immunofluorescence except for a zooming factor of 2.5 resulting in a pixel size of 0.248μm.

### Quantification of UNC-45A signal intensity in mouse and human tissues

The protocol to measure UNC-45A signals inside and outside areas of amyloid plaques was adopted from Sadleir et al. [12]. Briefly, the inside plaque area was defined by tracing the contour of an amyloid plaque detected by the amyloid β antibody with a free-handed selection tool (inside plaque ROI). The raw integrated density (Raw IntDen = sum of all pixel values) of UNC-45A signals within this area was measured and divided by its area to obtain the UNC-45A signal intensity within the inside plaque area. The outside plaque area contour was drawn approximately 30 μm away from the plaque contour (outside plaque ROI). The outside plaque area was calculated by subtracting the area of inside plaque ROI from the area of outside plaque ROI. Similarly, the UNC-45A Raw IntDen of the outside plaque area was calculated by subtracting UNC-45A Raw IntDen of the inside plaque ROI from the outside plaque ROI. The UNC-45A signal intensity outside plaque area was then obtained by dividing this value by the outside plaque area. All measurements were obtained from 20x images. All plaques greater than 600μm^2^ in mouse hippocampi and greater than 300μm^2^ in human hippocampi were quantified.

### Primary neuronal culture

C57BL/6J mouse (Jackson Laboratories, strain 000664) hippocampal neurons were cultured as previously described [64, 65]. Briefly, hippocampi from mouse pups at postnatal days 0-1 were removed, placed in a cold Hibernate-A medium (Brain Bits), then digested in a digestion medium containing Earle’s Balanced Salt Solution (EBSS) (Sigma) supplemented with 2g/L glucose, 0.5mg/mL papain, 0.2mg/mL L-cysteine, and 0.5mM ethylenediaminetetraacetic acid (EDTA) for 15 min at 37℃ with occasional gentle shaking. The hippocampi were gently washed in a digestion inhibitor solution containing EBSS supplemented with 2g/L glucose, 1mg/mL Bovine Serum Albumin (BSA), 1mg/mL trypsin inhibitor, and 1ug/mL DNase, then triturated. The cells were plated on poly-L-lysine coated coverslips in 24-well plates (30,000 cells/well) in a plating medium containing Minimum Essential Medium (MEM) supplemented with 10% heat-inactivated Fetal Bovine Serum (FBS), 5% heat-inactivated Horse Serum (HS), 200uM L-cystine, 0.5mg/mL penicillin/streptomycin/glutamine (PSG), and 1mM sodium pyruvate. At DIV2, cells were treated with 1uM AraC in a neurobasal medium supplemented with 5% HS, 0.5mg/mL PSG, 1mM sodium pyruvate, and 2% B27 for 8 hrs to inhibit glia proliferation. 80% of the neurobasal medium was changed to a fresh medium without AraC every 2-3 days.

### Neuroblastoma cell line

Undifferentiated N2A was cultured at 37℃ with 5% CO_2_ in Dulbecco’s Modified Eagle Medium (DMEM) with 10% FBS and plated on poly-L-lysine coated coverslips in 24-well plates or poly-L-lysine coated 24-well plates (30,000 cells/well for immunofluorescence staining, 50,000 cells/wells for drug treatments, and 75,000 cells/well for lentivirus infection). Undifferentiated N2A was differentiated by switching the medium to DMED with 0.5% FBS as previously described [66].

### Fixed cell immunofluorescence

Immunofluorescence staining of primary neurons and N2A was performed as described previously [12]. Briefly, cells were fixed for 20 min in 4% paraformaldehyde with 0.12M sucrose in PBS at 37℃, then permeabilized for 10 min in 0.5% Triton-X 100. Subsequently, the cells were blocked for 30 min in 5% BSA in PBST, stained with anti-UNC-45A (1:100 dilution in 2.5% BSA) for 1 hr, and anti-mouse Alexa 594 (1:250 dilution in 2.5% BSA) for 1 hr. The cells were mounted with Prolong Gold with DAPI (Thermo Fisher Scientific) and imaged with 20x/0.45 NA ELWD ADM S Plan FL objective and 40x/0.6 NA ELWD Plan FL DM objective (Nikon) on a Nikon TiE inverted microscope using Zyla 4.2 PLUS sCMOS camera (Andor) and NIS Element software (Nikon). Individual neurite swellings were imaged with a DIC microscopy on a Nikon TiE inverted microscope with 100x/1.49 NA Apo TIRF objective (Nikon). The minimum measurable distance for two pixels was 0.13μm. All images were obtained using the identical camera, microscope, and imaging criteria such as gain, brightness, contrast, and exposure time. Digital gray values of image pixels representing arbitrary fluorescence units (AFU) were obtained using Fiji software.

### Drug studies

To induce neurite swellings, differentiated N2A (DIV5) was treated with H_2_O_2_ and taxol and used concentrations varying 0uM (water) – 100uM and 0nM (DMSO) – 500nM for 1hr, respectively.

The determined optimum concentrations were used to treat GFP and UNC-45A-GFP expressing cells.

### Modulation of UNC-45A expression levels in cells

Lentiviral supernatant containing UNC-45A-GFP overexpression and GFP-empty vector control were prepared and used to infect differentiated N2A cells as previously described [30, 34].

### Western blot analysis

Primary neurons and N2A cells were lysed in 1% SDS lysis buffer. The total cellular protein (15µg) from each sample was separated by SDS-PAGE, transferred to PVDF membranes and subjected to western blot analysis using the specified antibodies. Amido Black staining was performed to confirm equal protein loading.

### Live cell imaging

The bright field images of cells were taken quickly at room temperature with a Nikon Eclipse TE200 fluorescent microscope using 20x/0.40 NA LWD and an NIS Element F3.3 camera and software (Nikon). The images were taken in areas with similar cell density across all conditions.

### Data analysis for neurite swelling

Neurite swelling sites were determined as previously described [43]. Briefly, the presence of swellings was determined by measuring the diameter of bulges along the length of an imaged neurite. The diameter of non-swelling sites was determined by averaging the diameter of adjacent neurite segments to swelling sites (1um away from the edge of the swelling). A bulge with a diameter greater than twice that of the adjacent neurite was considered a swelling (>2X). A bulge with a diameter less than twice that of the adjacent neurite was considered a swelling (<2X). For analysis of UNC-45A localization in neurite swelling, only >2x swellings were quantified. By using fixed area-circle ROI, UNC-45A gray value was measured at the center of the swelling site and non-swelling site 1um away from the edge of the swelling site. The circular ROI area was determined by the diameter of the non-swelling site. The same sized ROI was then moved to outside of neurites adjacent to the swelling or non-swelling sites and the local background was recorded. The background gray value was subtracted from the UNC-45A gray value to obtain UNC-45A AFU. For primary neurons, swelling sites on only thick neurites (>0.6um) were included in the quantification. For analysis of the number of swellings, the total number of swellings was counted manually within squire ROI (67134 μm^2^) and normalized by the total length of neurites within ROI. Neurite length was measured by the Fiji plugin NeuronJ as described previously [67].

### Statistical analysis for neurite swelling

Results are reported as mean±s.d. of three independent experiments unless otherwise indicated. The statistical significance of the difference was assessed with an unpaired two-tailed Student’s *t*-test using Prism (V.9 GraphPad) and Excel (Microsoft). The level of significance was set at *P*<0.05.

### Recombinant proteins

GFP(A206)-tagged human UNC-45A full-length (UNC-45A WT; 1–944 aa) was cloned, expressed, affinity purified, and dialyzed, as we previously described [31]. Recombinant GFP-human katanin p60 was purified as previously described [68].

### Preparation of microtubules

Porcine brain tubulin (T240), rhodamine-labeled tubulin (TL590M), and biotinylated tubulin (T333) were purchased from Cytoskeleton. Guanosine-5’-[(α,β)-methyleno] triphosphate (GMPCPP) was purchased from Jena Bioscience (NU-405S, Jena Bioscience). Single-cycled GMPCPP microtubules were made as previously described [31]. For microtubule binding assay with GFP-UNC-45A, these microtubules were treated with different concentrations of taxol (0, 10, 100μM) for 1hr or without/with 200μM H_2_O_2_ overnight at 37℃. For microtubule severing/depolymerization assay with GFP-UNC-45A, microtubules were treated with 10μM taxol for 1hr at 37℃. Double-cycled GMPCPP microtubules were made as previously described [24, 25]. The protocol for polarity-marked GMPCPP microtubules was adapted from a previously published method [69]. Specifically, 30 μl of elongation mixture, consisting of 0.5μM tubulin (30:1 mixture of unlabeled and rhodamine-labeled tubulin, respectively) and 0.5mM GMPCPP in BRB80 (80 mM 1,4-piperazinediethanesulfonic acid, 1 mM MgCl2, 1 mM EGTA, pH 6.9—pH was adjusted with KOH), were kept on ice for 5 mins, before warming up to 37℃ for 1 min. 3μL of 2μM GMPCPP seeds were then added to the elongation mixture and incubated at 37 °C for 30 mins. The mixture was replenished with 0.5μM tubulin mixture every 30 mins for 1 hr and 30 mins. The elongated microtubules were then diluted 60x in warm BRB80 containing 10μM taxol in order not to break microtubules and reduce the concentration of free tubulin, then introduced into a TIRF chamber. The polarity-marked microtubules were incubated in the chamber at room temperature for 15 mins, washed with warm BRB80 containing 10μM taxol and 1mM DTT three times to remove the remaining free tubulin, and incubated with 10μM taxol for at least 30 mins at 37 °C before imaging. Taxol-stabilized GTP microtubules were prepared as previously described [31, 70].

### Microtubule binding/ depolymerization/ severing assay

TIRF chambers were prepared as we previously described [47]. 15-20 single-cycled GMPCPP microtubules/ field of view or 5 polarity-marked GMPCPP microtubules were introduced into the TIRF chamber and washed with 4 channel volumes of warm BRB80 containing 1mM DTT. For the binding/depolymerization assays, 250nM GFP-UNC-45A in assay solution (50 mM KCl, 0.5% Pluronic F127, 0.2 mg/ml casein, 1.5% glycerol, 0.1% methylcellulose 4000 cP, 20 mM glucose, 110 μg/ml glucose oxidase, and 20 μg/ml catalase, 20 mM DTT in BRB80) was introduced as previously described [47]. For the severing assay, the assay buffer containing 0.2mg/mL casein, 20mM glucose, 110μg/mL glucose oxidase, 60μg/mL catalase, and 10mM DTT in BRB80 was introduced with various concentrations of GF-UNC-45A. The images for the binding assay were taken at 5 min after the addition of GFP-UNC-45A. For depolymerization and severing assays, the video recording was started immediately after the addition of GFP-UNC-45A, and recorded for 20 min. The binding assay of 25nM GFP-katanin to taxol GTP microtubules was performed as previously described [71] with 2.5% glycerol in the assay buffer. All experiments were performed at 37℃.

### Microtubule repair assay

For the repairment of rhodamine-labeled microtubules with HiLyte 488-tubulins (Cytoskeleton, TL488M), 15-20 double-cycled GMPCPP microtubules / field of view were introduced into the TIRF chamber, treated with either taxol (0, 10, 100μM) for 1hr or H_2_O_2_ (0, 0.25, 0.5, 1mM) for 40 mins at 37℃. The protocol for tubulin repair assay was adapted from a previously published protocol [24]. Briefly, the chamber was washed with 10 channel volumes of a pre-perfusion solution containing 2.5mg/mL casein, and 5% Pluronic F127 in BRB80. Tubulin aggregates were ultracentrifuged at 279,000×g for 10 min at 4℃ and tubulin concentration was measured by A280 prior to each experiment. Tubulin solution containing 0.5mM GTP, 1% Pluronic F127, 2.5mg/mL casein, and 500nM HiLyte 488-tubulins was introduced into the chamber and incubated for 5 min, washed out with 10 channel volumes of tubulin wash solution containing 1.5mg/mL casein and 1% Pluronic F127 in BRB80, and imaged with tubulin imaging solution containing 1.5mg/mL casein, 1% Pluronic F127, 20mM glucose, 110μg/mL glucose oxidase, 60μg/mL catalase, and 10mM DTT in BRB80.

For the repairment of biotin-labeled microtubules with rhodamine-tubulins in the presence of GFP-UNC-45A, less than 5 double-cycled GMPCPP microtubules /field of view were introduced into the TIRF chamber, treated with 100μM taxol for 1hr at 37℃. The chamber was washed with 10 channel volumes of a pre-perfusion solution, 250nM GFP-UNC-45A and rhodamine-tubulins (concentrations varying from 250nM to 5μM) in UNC-45A binding solution containing 1mM GTP, 2.5mg/mL casein, 1% Pluronic F127, 20mM glucose, 110μg/mL glucose oxidase, 60μg/mL catalase, and 10mM DTT in BRB80 was introduced into the TIRF chamber, incubated for 8 min, and washed out with 10 channel volumes of UNC-45A wash solution containing 2.5mg/mL casein, 5% Pluronic F127, 1% tween 20 in 20mM HEPES at pH7.5. Then the tubulin imaging solution was introduced to the chamber, and images were taken.

### Total internal fluorescence (TIRF) Imaging

For two-color TIRF imaging, time-lapse videos and fluorescent images of rhodamine-microtubules and GFP-UNC-45A, GFP-katanin, or HiLyte 488-tubulins were obtained with 488 nm and 561 nm lasers generated from TIRF mode on a Zeiss Axio observer Z1 inverted microscope using Alpha Plan-Apochromat 100×/1.46 NA objective lens as previously described [31]. The standard exposure time was 20ms for both 488 nm and 561 nm lasers, and 100ms exposure was used for HiLyte 488-tubulins. All experiments were done at 37℃. For two-color TIRF with differential interference contrast (DIC) microscope, images of biotin-labeled microtubules with rhodamine-tubulins and GFP-UNC-45A were obtained with 488 nm and 561 nm TIRF lasers generated from Nikon TiE inverted stand using a 100x/1.49NA Apo TIRF objective lens. The exposure time was 200ms for both 488 nm and 561 nm lasers. All experiments were done at 37°C using an objective heater (OkoLab).

### Co-sedimentation of free tubulins and GFP-UNC-45A

Pre-spun aggregate-free GFP-UNC-45A and free tubulins were mixed at final concentrations of 250nM and 5μM, respectively in BRB 80 buffer with 1mM GTP on ice. The mixture was quickly warmed up to 37℃, incubated for 8 min, and ultracentrifuged at 20K, 40K, or 85K rpm at 4℃ for 10 min with Optima TLX ultracentrifuge TLA-100 rotor (Beckman). The supernatant and pellet were carefully separated, and the pellet was resuspended in the original mixture volume of BRB80. 18μL of proteins were boiled with 4% SDS and 1x laemmli buffer, ran on a SDS-PAGE gel, and stained with Coomassie blue.

### Data analysis for *in vitro* microtubule experiments

For the measurement of fluorescence intensity of GFP-UNC-45A, HiLyte 488-tubulins, and rhodamine-tubulins on microtubules, the protocol was adapted from previously published work [72]. Fiji was used to draw a rectangle ROI covering the entire length of the microtubule of interest with a fixed width and gray value was measured on GFP-UNC-45A, HiLyte 488-tubulins, or rhodamine-tubulins channel. The same-sized ROI was then moved adjacent to the microtubule and the local background was recorded. Measured gay values were corrected to the length of the microtubules. The background value was then subtracted from the value of interest to give a corrected intensity measurement. Finally, the average background-corrected intensity along microtubules in no treatment group was normalized to 1, and the average background-corrected intensity in treatment groups was normalized to no treatment group. To measure the GFP-UNC-45A signal at microtubule crossing and non-crossing sites, a fixed-sized circular ROI covering the entire width of microtubule at a crossing and non-crossing sites 2μm away from the crossing site, and the integrated density from the ROI was measured on the GFP-UNC-45A channel. To measure the GFP-UNC-45A signal at microtubule bundling and non-bundling segments, a rectangle ROI covering the entire segment of the microtubule bundling of interest with a fixed width and gray value was measured on GFP-UNC-45A. The measured gray values were corrected to the length of the bundling and background values as described above. The average background-corrected intensity of non-crossing and non-bundling groups was normalized to 1, and the crossing and bundling groups’ intensity was normalized to non-crossing and non-bundling groups. The duration of association, binding frequency, severing frequency, severing %, and depolymerization rate were measured and calculated as previously described [52, 73].

### Statistical analysis for *in vitro* microtubule experiments

Results are reported as mean±s.d. of three or more independent experiments. Unless otherwise indicated, the statistical significance of the difference was assessed with an unpaired two-tailed Student’s *t*-test using Prism (V.9 GraphPad) and Excel (Microsoft). The level of significance was set at *P*<0.05.

## Acknowledgements

We gratefully acknowledge Nimisha Krishnan and Jennifer L. Ross (Department of Physics, Syracuse University, Syracuse, NY, USA) for their generous gift of the katanin plasmid used for the preparation and purification of recombinant katanin.

