## Supplementary for "UNC-45A drives ATP-independent microtubule severing via defect recognition and repair inhibition, contributing to neurite dystrophy"

Supporting Figure 1

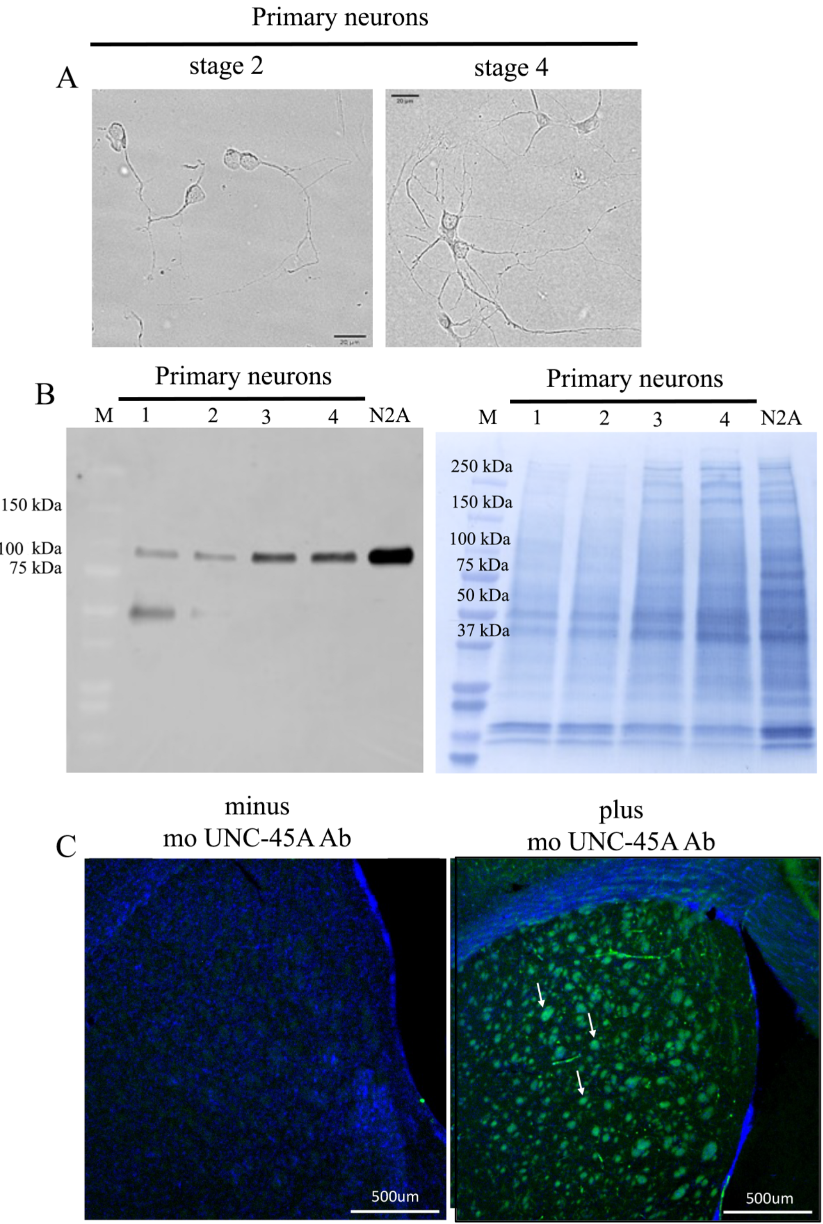

### Supporting Figure 2

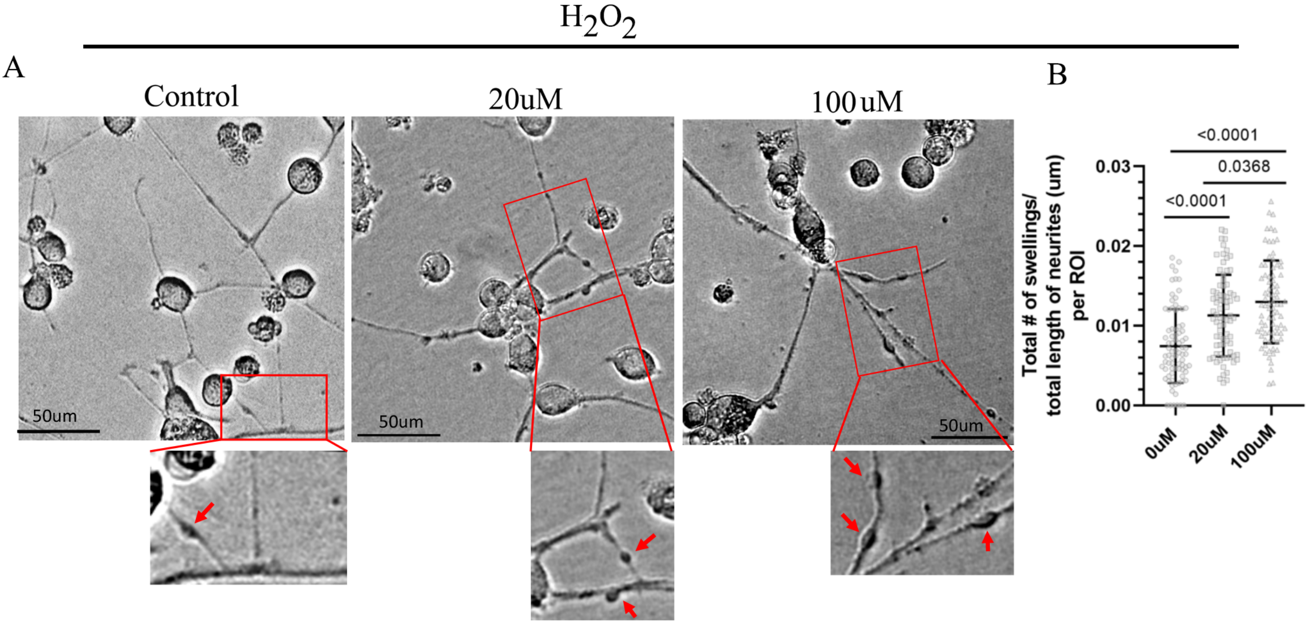

Supporting Figure 3

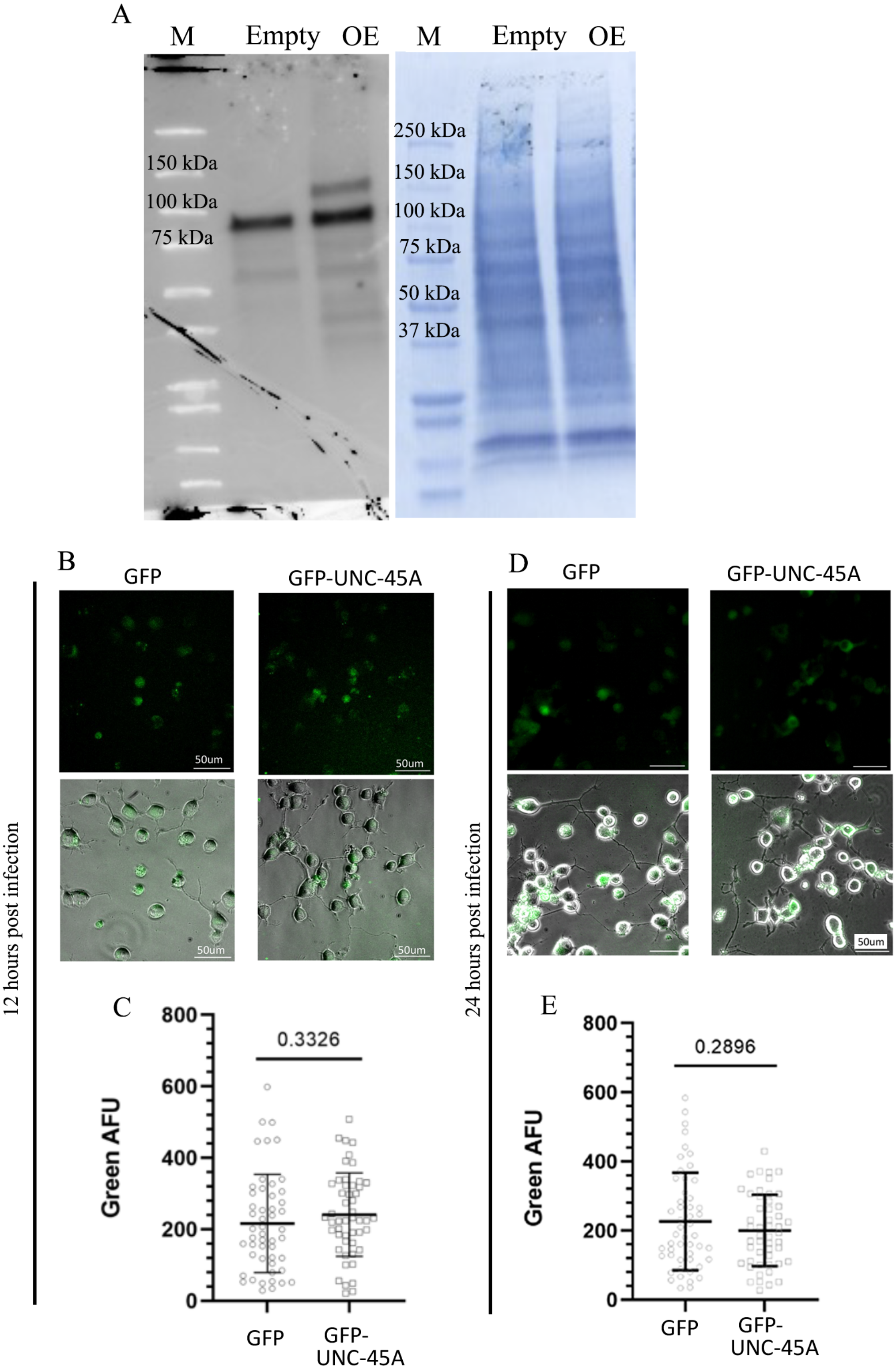

Supporting Figure 4

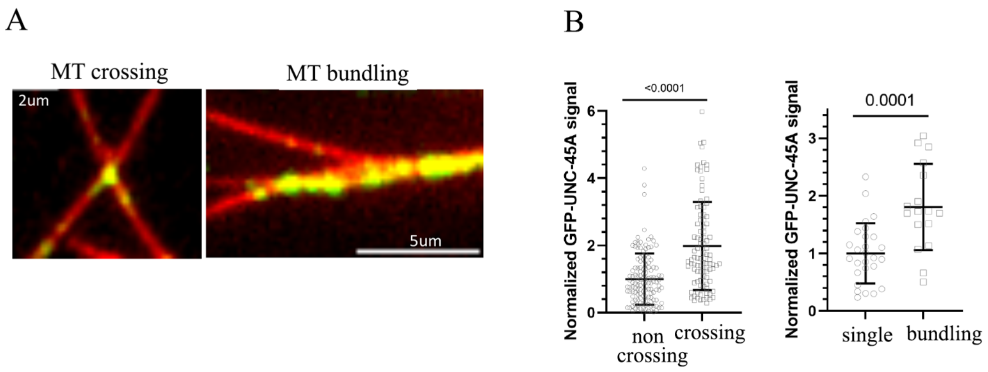

Supporting Figure 5

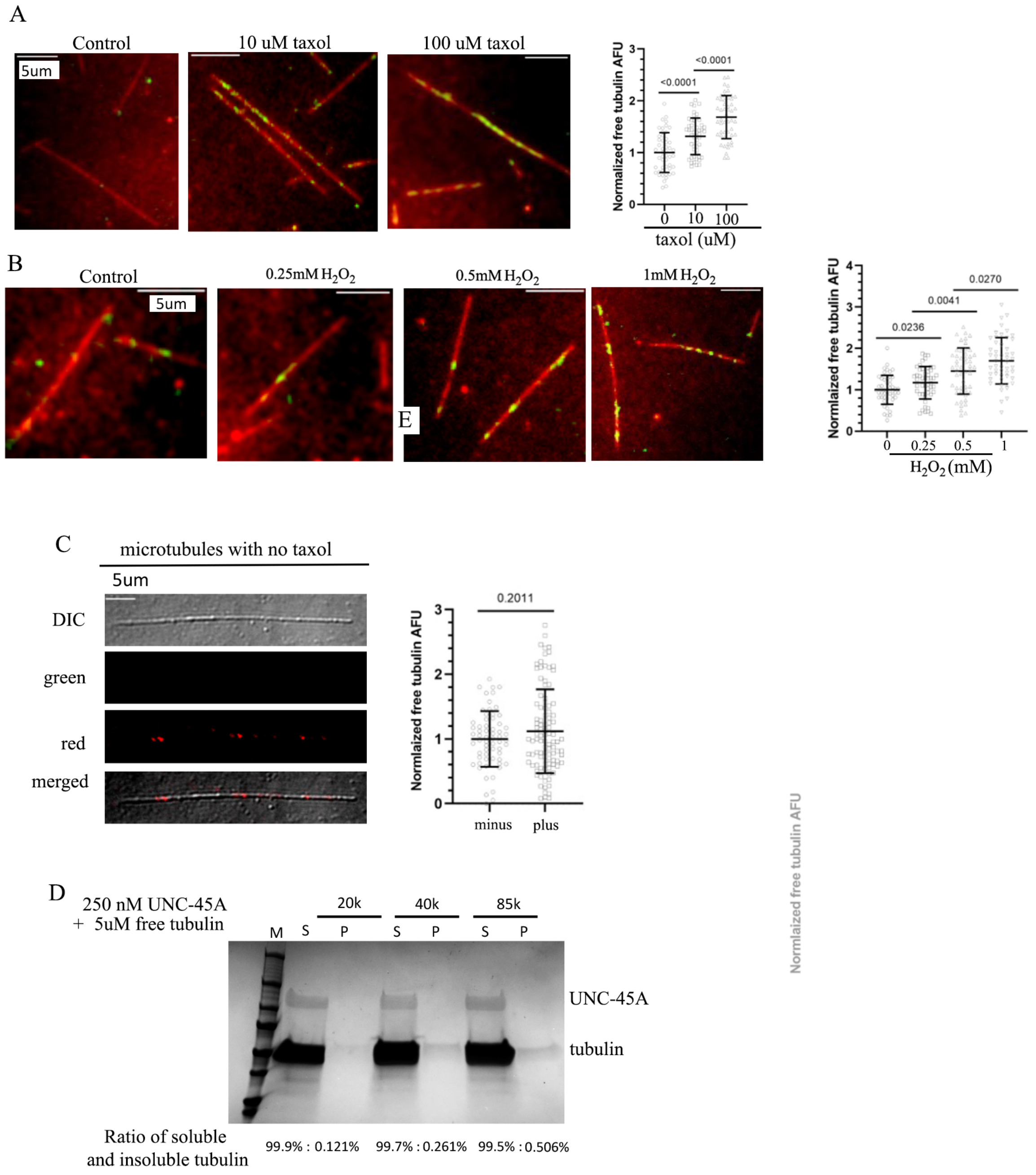

**Extended Data Figure Legends**

**Extended Data Figure 1**. **Mouse monoclonal UNC-45A antibody specifically recognizes UNC-45A**. **A**. Representative images of mouse primary hippocampal neurons at stage 2 (DIV1) and stage 4 (DIV5) harvested for western blot analysis. **B**. *Left*, Western blot analysis with cell lysates from mouse primary hippocampal neurons at different developmental stages. *Lane 1*= stage 2 (DIV1). *Lane 2*= stage 3 (DIV2). *Lane 3*= stage 4 (DIV5). *Lane 4*= stage 5 (DIV14). Lysate from the N2A neuroblastoma cell line was used as positive control. M= marker. *Right*, total protein loading was assessed by amido black staining. C. Immunofluorescence analysis of non-transgenic mouse brain (striatum) stained (plus) or not (minus) with anti-UNC-45A antibody followed by Alexa 488 secondary antibody (green). Blue=DAPI. White arrows indicate axons rich areas. The images were acquired with identical exposure with a 4x objective.

**Extended Data Figure 2. UNC-45A overexpression in differentiated N2A. A.** *Left*, Western blot analysis showing the levels of UNC-45A in differentiated N2A expressing GFP-empty vector (Empty) and UNC-45A-GFP (OE) at 12 hours of post-infection. *Right*, amido black was used as a loading control. M=marker. **B**. Representative images of GFP or UNC-45A-GFP expressing differentiated N2A cells (at 12 hours of post-infection) having similar amounts of GFP. *Top* =Green channel. *Bottom* = Green channel merged with phase contrast images. **C**. Quantification of GFP (AFUs) in GFP and UNC-45A-GFP expressing cells shown in B (mean±S.D., n = 50 cells per condition). **D**. Representative images of GFP or UNC-45A-GFP expressing differentiated N2A cells (at 24 hours of post-infection) having similar amounts of GFP. *Top* =Green channel. *Bottom* = Green channel merged with phase contrast images. **E**. Quantification of GFP (AFUs) in GFP and UNC-45A-GFP expressing cells shown in D (mean±S.D., n = 50 cells per condition).

**Extended Data Figure 3. $H_2O_2$ treatment result in a dose-dependent increase in swellings in the neurites of differentiated N2A cells. A.** *Top*, Representative images of differentiated (DIV5) N2A cells treated with vehicle (water) or $H_2O_2$ at the indicated concentrations for 1 hour. Bright-field images were acquired with a 20x objective. *Bottom*, close-up images of neurite swellings shown in the *top panels*. Red arrows indicate the sites of neurite swellings. **B**. Quantification of neurite swellings shown per each condition A (mean±S.D., n= 80 ROIs per condition).

**Extended Data Figure 4. UNC-45A preferentially binds to crossing and bundling regions of the microtubules. A**. *Left*, representative image of crossing site of rhodamine-labeled GMPCPP microtubules (red) and 250nM GFP-UNC-45A (green). *Right*, representative image of bundling site of rhodamine-labeled GMPCPP microtubules (red) and 250nM GFP-UNC-45A (green). **B**. *Left*, Quantification of GFP-UNC-45A binding to non-crossing or crossing microtubules. The fluorescent intensity of GFP-UNC-45A at crossing sites and sites 2 μm away from the crossing sites were quantified (mean±S.D., n= 132 non-crossing sites, and 97 crossing sites). *Right*, Quantification of UNC-45A binding to single versus bundling microtubules. The fluorescence intensity of GFP-UNC-45A on microtubule bundling segments and single microtubules segments were quantified (mean±S.D., n= 27 single segments and 17 bundling segments).

**Extended Data Figure 5. In a dynamic system, damaged microtubules are repaired via incorporation of free tubulin. A.** *Left*, representative images of rhodamine-labeled MTs (red) treated with different concentrations of taxol (0, 10, 100μM) for 1 hr and repaired with 500nM

HyLight 488-tubulins (green). *Right*, quantification of fluorescent intensity of HyLight 488-tubulins binding to microtubules and expressed as normalized free tubulin AFU. n= 50 MTs per condition were quantified. Error bars represent mean ± SD. **B**. *Left*, representative images of rhodamine-labeled MTs (red) treated with different concentrations of $H_2O_2$ (0, 0.25, 0.5, 1mM) for 40 min and repaired with 500nM HyLight 488-tubulins (green). *Right*, quantification of fluorescent intensity of HyLight 488-tubulins binding to microtubules expressed as normalized free tubulin AFU. n= 50 MTs per condition were quantified. Error bars represent mean ± SD. **C**. Coomassie staining of tubulin and GFP–UNC-45A at the indicated concentrations, following centrifugation at 20,000, 40,000, or 85,000 rpm. Both supernatant and pellet fractions were loaded onto the gel. M = protein marker. **C.** *Left***,** representative images of non taxol-treated biotinylated microtubule repaired with 500nM rhodamine-tubulins (red) in the absence of GFP-UNC-45A (green). *Right*, quantification of fluorescent intensity of rhodamine-tubulins binding to undamaged microtubules in the absence of GFP-UNC-45A and to damaged microtubules by 100µM taxol in the presence of 100nM GFP-UNC-45A. n= 100 microtubules per condition were quantified. Error bars represent mean ± SE. **D.** Pre-spun, aggregate-free GFP–UNC-45A (250 nM) was mixed with free tubulin (5 µM) and subjected to ultracentrifugation at 20,000, 40,000, or 85,000 rpm. Supernatant (S) and pellet (P) fractions were separated and analyzed by Coomassie blue staining to detect GFP–UNC-45A and tubulin distribution.
